# Hydrogen-deuterium exchange coupled to top- and middle-down mass spectrometry enables high-resolution measurements of histone tail dynamics before and after nucleosome assembly

**DOI:** 10.1101/310177

**Authors:** Kelly R. Karch, Mariel Coradin, Levani Zandarashvili, Zhong-Yuan Kan, Morgan Gerace, S. Walter Englander, Ben E. Black, Benjamin A. Garcia

## Abstract

Until recently, a major limitation of hydrogen deuterium exchange mass spectrometry (HDX-MS) was that resolution of deuterium localization information was limited to the length of the peptide generated during proteolysis. Recently, however, it has been demonstrated that electron transfer dissociation (ETD) allows for preservation of deuterium label in the gas phase and therefore can be used to obtain more resolved information. To date, this technology has remained mostly limited to single, small, already well-characterized model proteins. Here, we optimize, expand, and adapt HDX-MS/MS capabilities to accommodate histone and nucleosomal complexes on top-down (TD) HDX-MS/MS and middle-down (MD) HDX-MS/MS platforms and demonstrate that near site-specific resolution of deuterium localization can be obtained with high reproducibility. We are able to study histone tail dynamics in unprecedented detail, which have evaded rigorous analysis by traditional structural biology techniques for decades, revealing important novel insights into chromatin biology. This work represents the first heterogeneous protein complex and protein-DNA complex to be analyzed by TD- and MD-HDX-MS/MS, respectively. Together, the results of these studies highlight the versatility, reliability, and reproducibility of ETD-based HDX-MS/MS methodology to interrogate large protein and protein/DNA complexes.

## Introduction

Hydrogen/deuterium exchange coupled to mass spectrometry (HDX-MS) is a technique used to monitor the structure and dynamics of proteins in solution. In an HDX-MS experiment, natively folded proteins are diluted into D_2_O, allowing accessible amide protons to be exchanged for deuterium atoms in the solvent. The rate of deuterium exchange acts as a proxy for protein structure and stability, with a more stable structure experiencing a slower rate of exchange (and hence higher degree of protection from exchange) than a less stable structure. Given that protein dynamics are closely tied to function, HDX-MS has proven to be a valuable tool in a wide range of applications such as monitoring the effects of post-translational modifications, mapping protein interactions, and measuring changes in structure and stability upon specific perturbations (i.e. mutations, truncations).

Most HDX-MS studies to date incorporate bottom-up mass spectrometry (BU-HDX-MS), whereby proteins are digested (usually with pepsin) into peptides, and the deuterium content of each peptide is measured (Figure 1). This technique allows the user to obtain information at the resolution of the length of the peptide (generally 5-20 amino acids). If many overlapping peptides are obtained, the user can potentially obtain more site-specific information by subtracting the deuterium content of overlapping peptides.^1–4^ Therefore, the degree of resolution of deuterium localization obtained in BU-HDX-MS is dependent upon how many overlapping peptides are detected, with a larger number of overlapping peptides providing.

**Figure 1.**
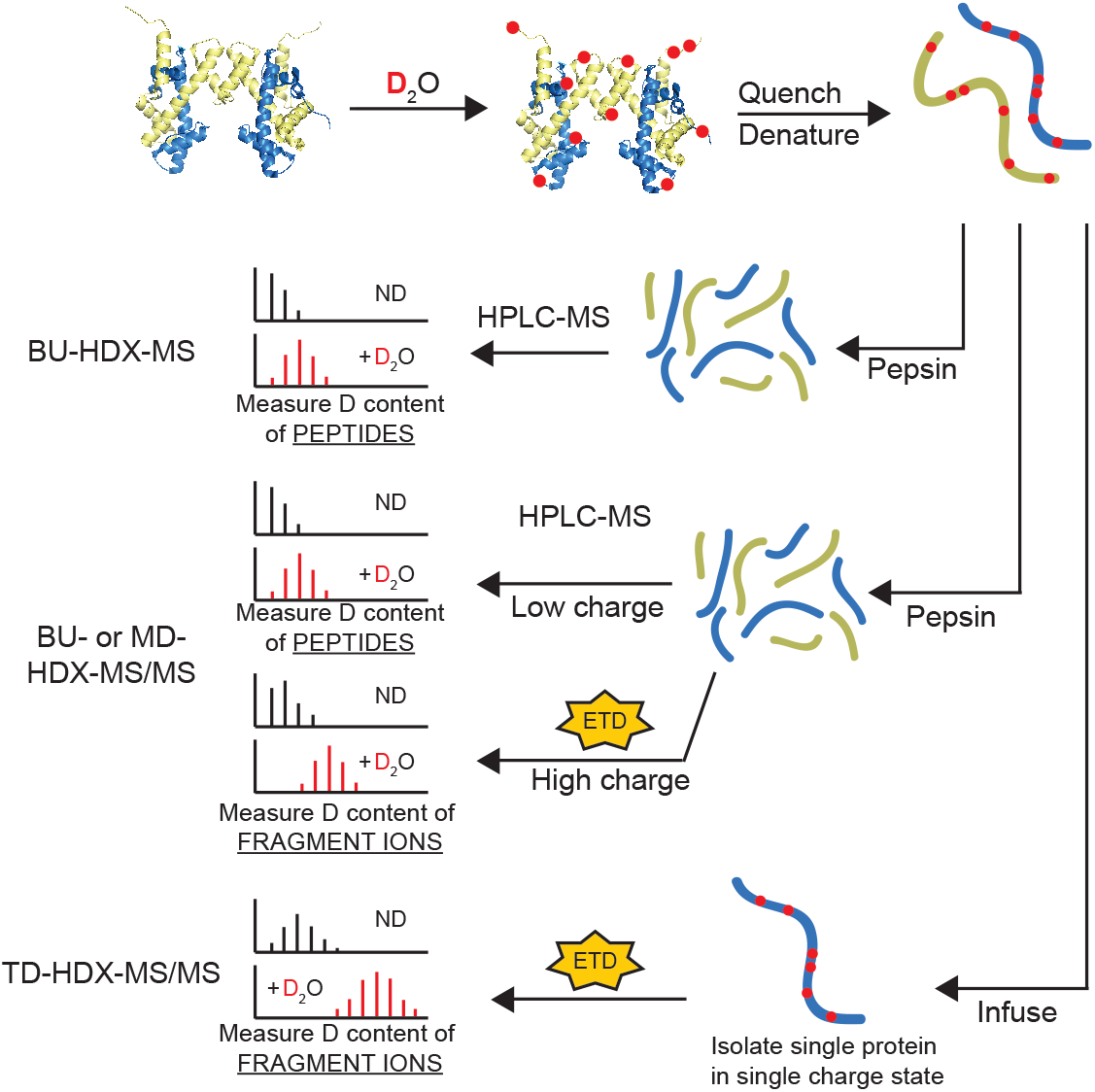
Schematic of HDX-MS Experiments. Proteins are diluted into D_2_O buffer and allowed to exchange over time. Aliquots are quenched at different timepoints and deuterium content is measured by mass spectrometry. In BU-HDX-MS experiments, proteins are digested with pepsin, separated by RP-HPLC, and the deuterium content of the intact peptides are measured. In BU- or MD-HDX-MS/MS experiments, the same sample processing occurs but highly charged peptides can be targeted for fragmentation to obtain more resolved information. BU-HDX-MS/MS is used for analysis of standard sized peptides, and MD-HDX-MS/MS refers to long peptides (generally > 20 amino acids). In TD-HDX-MS/MS experiments, intact proteins are infused into the mass spectrometer and fragmented by ETD. Deuterium content of fragment ions are measured. ND = non-deuterated.

Fragmenting peptides in an MS/MS experiment provides the opportunity to obtain more resolved information. In this process, peptide ions are fragmented inside the mass spectrometer and the masses of the fragment ions are measured, allowing for deuterium content to localized to specific residue(s). Up to site-specific deuterium content measurements can be obtained depending on the efficiency of fragmentation. However, one barrier to using fragmentation in HDX-MS/MS experiments is the phenomenon of “scrambling,” whereby amide protons and deuterium atoms migrate along the backbone in the gas phase upon high vibrational energy. Scrambling has precluded collision-based fragmentation methods from HDX studies due to the high vibrational energy imparted to the peptide ions in the gas phase. However, electron transfer dissociation (ETD) fragmentation has been shown to preserve the deuterium label of ions and allow for accurate measurements of the deuterium content of fragment ions.^5–9^

HDX-MS/MS has been applied in both bottom-up (BU) and top-down (TD) mass spectrometry experiments. In BU-HDX-MS/MS, any ETD-amenable peptides are targeted for fragmentation, enabling more resolved information to be obtained for a portion of the protein. In TD-HDX-MS/MS, intact proteins are infused and fragmented, usually bypassing proteolysis and separation steps (Figure 1). TD-HDX-MS/MS has several advantages over BU-HDX-MS/MS, namely that full coverage of the protein is guaranteed and high-resolution information can potentially be obtained across the entire sequence.

Despite the great utility of HDX coupled to ETD MS/MS, very few studies have utilized these platforms to monitor protein structure and function. BU-HDX-MS/MS has been accomplished in a handful of studies, all of which have focused on a single protein or oligomers of a single protein.^5,10–14^ TD-HDX-MS/MS has been slightly larger in scope but has been mainly limited to small proteins (< 200 amino acids) that easily fragment, with a notable exception of the Borchers group who used TD-HDX-MS/MS to analyze antibodies.^6,13,15–25^ Together, these initial studies demonstrate the great power of ETD-HDX-MS/MS techniques to measure protein structure and stability in great detail and also highlight the potential of these methods to be applied to more complex samples.

Here, we sought to optimize, validate, and expand the capabilities HDX-MS/MS platforms using histone protein complexes. Histone proteins make a great case study for these aims as they can exist in a variety of complexes, are amenable to ETD fragmentation, have significant biological importance, and are difficult to study by traditional BU-HDX-MS.

Histone proteins are responsible for organizing DNA into higher order chromatin structure.^26,27^ They bind DNA in an octameric form containing two copies of each core histone: H2A, H2B, H3, and H4. This protein-DNA complex, called a nucleosome, binds approximately 147 base pairs of DNA.^28^ Histones can also exist in smaller complexes, namely an H2A/H2B heterodimer, and an (H3/H4)_2_ heterotetramer. Each histone protein contains an N-terminal tail that protrudes from the nucleosomal surface. These tails are critical for nucleosome function, including transcription and formation of higher order structure. Whether the tails have structure in solution and how the dynamic properties of the histone tail affects function have been outstanding questions in the chromatin field for decades.^29,30^ Answering these questions will shed light into basic nuclear processes and may also aid in the development of therapeutics as aberrant chromatin structure is tightly linked to many diseases, including cancer. Currently, there is no rigorous method to study histone tail structure and stability as nucleosomes are too large for some techniques such as NMR, and the flexibility of the N-terminal tails makes it difficult to crystalize or simulate in molecular dynamics approaches. Previous studies have utilized traditional BU-HDX-MS methodology (without ETD), but the tail domains were either not detected or were very long (~50 amino acids) and so highly resolved information could not be obtained.^31,32^

In this study, we performed TD-HDX-MS/MS on histone monomers and tetramers, representing the first protein complex to be analyzed with this method. We also developed a middle-down (MD) HDX-MS/MS platform whereby proteins are digested and long peptides (in our case, 39 to 49 amino acids long) are fragmented with ETD. We used this platform to study histone tetramers and nucleosomes, representing the first protein/DNA complex to be analyzed by HDX-MS/MS. In doing so, we were able to rigorously address histone tail structure and dynamics in nucleosomal and sub-nucleosomal complexes, which has evaded in-depth analysis by traditional methods for decades. We were able to obtain highly resolved, reproducible, and comparable deuterium content measurements on both platforms, highlighting the strength of these methods to reliably study protein dynamics in great detail. These methods can easily accommodate nearly any protein or protein complex of interest, demonstrating the adaptability and broad utility of HDX-MS/MS.

## Materials and Methods

### Protein expression, purification, and reconstitution

Human histones H2A, H2B, H3, and H4 were expressed in BL21 [DE3] (pLysS) cells and purified as monomers as previously described.^33,34^ Briefly, protein was extracted from cells and separated by gel filtration (column: HP Sephacryl 26/60 S200). Fractions containing the histone of interest were pooled, dialyzed into cation exchange buffer, and purified by cation exchange (column: HiTrap 5mL SPFF). Histone fractions were pooled, dialyzed into low salt buffer, and lyophilized for long-term storage. Histone H2A was expressed with a His-tag at the N-terminus, and an additional purification step with a nickel column was performed prior to gel filtration. The tag was cleaved with PreScission protease, leaving a small amino acid tag at the N-terminus with the sequence GPLG.

Histones were refolded into H2A/H2B dimers and (H3/H4)_2_ tetramers by mixing equimolar amounts of the constituent proteins followed by dialysis as previously described.^28,34^ Briefly, histone proteins were resuspended in urea buffer. Urea was slowly dialyzed out, and the resulting complex was purified using gel filtration and concentrated to approximately 1ug/uL.

DNA, consisting of the 195 base pair 601 positioning sequence, was made using PCR from a plasmid containing the sequence of interest. The PCR reactions were pooled, DNA was precipitated with ethanol and purified by cation exchange chromatography.^35^ Fractions containing the DNA of interest were pooled, precipitated and resuspended to an appropriate concentration for use in nucleosome reconstitution.

Nucleosomes were reconstituted as previously described by combining the components in a 1:2:1 molar ratio of DNA, H2A/H2B dimers, and (H3/H4)_2_ tetramers. Slow dialysis into low salt buffer allowed for formation of nucleosome particles, which were validated by native gel electrophoresis.^33,34^

### Top-Down HDX/MS

#### Scrambling analysis

We used peptide probe ‘P1’ (AnaSpec, Inc.) to determine the degree of scrambling as described.^8^ Briefly, the powdered peptide was dissolved in 99.9% D_2_O (Sigma) to a concentration of 1 ug/uL and incubated at room temperature for at least 24 hours. Peptides were diluted 50-fold into quench buffer (50% acetonitrile, 0.1% formic acid, pH 2.5 in H_2_O) for 10 seconds followed by freezing on dry ice. Samples were thawed and infused into the mass spectrometer as described below and fragmented with ETD using the following parameters: isolation window: 10m/z, S lens RF: 60%, ETD reaction time: 70ms, resolution: 60,000, capillary temperature: 150°C. Data was analyzed as previously described,^9^ except that centroids were calculated using ExMS2 software from the Englander lab.^36^

#### HDX and sample preparation

Lyophilized monomeric H4 proteins were resuspended in buffer (10mM Tris, 0.3mM EDTA in H_2_O, pH 7.0) to a final concentration of approximately 1ug/uL. Tetramer was also analyzed at this concentration in the same buffer. Deuterium exchange was conducted by mixing 20uL of protein (20ug) with 60uL of on-exchange buffer (10mM Tris, 0.3mM EDTA in D_2_O, pD 7.51) for the indicated time at 4°C. The exchange reaction was quenched by the addition of 120uL quench buffer (0.8% formic acid). Samples were immediately desalted using home-made C8 stage tip columns as previously described, except that the wash buffer consisted of 0.8% formic acid in dH_2_O, pH 2.25 and the elution buffer consisted of 75% acetonitrile/25% wash buffer.^37,38^ All steps in the desalting process were precisely timed to prevent differences in back exchange between samples. Before addition of the sample, the spin columns were activated with 75uL of methanol and washed with 75uL of quench buffer. The desalting steps are as follows: 1. Add sample, spin at 7000xg for 2 minutes; 2. Add 75uL of wash buffer, spin at 7000xg for 50 seconds; 3. Switch collection tube to a new clean tube, 4.

Add 20uL elution buffer, spin at 2400xg for 1 minute. Samples were transferred to a new precooled tube and immediately flash frozen in liquid nitrogen.

#### Infusion into MS

Samples were thawed in an ice water bath and immediately loaded into the pre-cooled (4°C) sample tray of an Advion Triversa Nanomate. The sample was delivered to a chip containing electrospray nozzles and infused. The spray quality was optimized in the first 30 seconds of infusion by altering the spray voltage (1.6 to 2.2 V) or air pressure of the Nanomate (0.4 to 0.5 psi) if needed. To prevent back exchange, a cooling device was used to lower the temperature of the sample as it infused into the mass spectrometer as described by the Jørgensen group.^39^ Briefly, we coiled approximately 60 feet of copper tubing inside an insulated box. The box was filled with dry ice, and nitrogen gas was flowed through the tubing at 25 psi and aimed directly on top of the infusion tip carrying the sample. The position of the gas nozzle was aligned perpendicular to the spray tip to minimize interference with sample spray. The temperature of the nitrogen gas exiting the cooling apparatus reached approximately -10°C.

#### Instrument Method

The capillary temperature was set to 150°C. All data were acquired manually. A full MS scan was acquired in the Orbitrap for approximately 30 seconds (resolution: 60,000; Scan range: 500-1300 m/z; S-lens RF level: 60%; AGC target: 5.0e5; maximum injection time: 100ms; source fragmentation: 35eV; 1 microscan). During this time, the centroid of the intact H4 in +15 charge state was approximated by eye. ETD MS/MS scans were then acquired in the Orbitrap (resolution: 120,000; scan range: 150-2,000 m/z; S-lens RF level: 60%; AGC target: 5.0e4; maximum injection time: 100ms; source fragmentation: 35 eV). The approximate centroid was specified for the precursor m/z. Ions were isolated in the quadrupole (isolation window: 8m/z) and fragmented with ETD (10ms) for approximately 3 minutes.

#### Data Analysis

HDExaminer (Sierra Analytics, version 2.5) was used to analyze deuterium content of the fragment ions, allowing +1 to +15 charge states. The intact sequence of H4 was imported into the software. Raw MS files were uploaded, and data was analyzed from minutes 1 to 2 of the data acquisition. All identified fragment ions were manually validated.

### Bottom-Up/Middle-Down HDX/MS

#### HDX

Deuterium exchange reactions were carried out on ice by mixing 5uL of protein (5ug) with 15uL of on-exchange buffer (10mM Tris, 0.3mM EDTA in D_2_O, pD 7.51). Reactions were quenched at the indicated time points by addition of 30 uL quench buffer (2.5M gdmCl, 0.8% formic acid, 10% glycerol in dH_2_O). Samples were immediately flash frozen in liquid nitrogen and stored at -80°C until MS analysis.

#### LC-MS/MS

Samples were thawed in ice water and injected into a cooled online sample processing system (3-6°C) composed of a pepsin column, a C18 trap column, and a C18 analytical column. A Shimadzu LC-10AD pump was used to pump the sample through an immobilized pepsin column at 0.05 mL/min onto a C18 trap column (1x 5mm, C18 PepMap100, Thermo Scientific, P/N 160434). Pepsin was immobilized by coupling to Poros 20 AL support (Applied Biosystems) and packed into column housings of 2 mm x 2 cm (64 ml) (Upchurch). Peptides were eluted onto and separated by an analytical C18 column (50 x 0.3mm, Targa 3um C18 resin, Higgins Analytical, Serial No. 269232) by a reverse-phase gradient delivered by an Agilent 1100 Series pump at 6uL/min (Buffer A: 0.1% formic acid, 0.05% TFA in H_2_O, pH 2.25 at room temperature; Buffer B: 0.1% formic acid in acetonitrile). The gradient consists of the following steps: 10-55% B in 15 minutes, 55-95% B in 5 minutes. The column was washed with 95%B for 30 minutes followed by re-equilibration at 10% B for 10 minutes between runs.

The sample was sprayed into a Thermo Orbitrap Fusion mass spectrometer. The capillary temperature was set to 215°C. For each construct, at least three non-deuterated samples were analyzed using HCD and ETD to identify peptides for data analysis. For the HCD method, a full MS scan was collected in the Orbitrap (resolution: 60,000; 360-1000 m/z, AGC target: 5 x 10^5^, maximum injection time: 50ms, RF lens: 55%) followed by a series of MS/MS scans for two seconds where ions are chosen for fragmentation sequentially based on their abundance. Fragment ions were measured in the ion trap (HCD collision energy: 30%, stepped collision energy: 5%, scan rate: rapid, maximum injection time: 200ms, AGC target: 1 x 10^4^, centroid mode). For the ETD method, a full MS scan was collected in the Orbitrap (same settings as HCD method), followed by ETD MS/MS scans based on abundance for two seconds in the Orbitrap (charge states: 5-10, ETD reaction time: 20 ms, ETD reagent target: 2 x 10^5^, resolution: 60,000, maximum injection time: 400ms, AGC target: 2 x 10^5^, 3 microscans). Tail peptides were targeted for fragmentation during their elution times (sequences in Table 4.3, target masses and charges in Table 4.4).

Given that HCD fragmentation leads to scrambling, all deuterated samples were analyzed using only ETD fragmentation. The same ETD MS method was used as specified for the non-deuterated samples; however, different masses, corresponding to the approximate centroid of the isotopic distribution, were targeted according to Table 4.4 during their respective elution times. In tetramer runs, only H3 and H4 tails were targeted. In nucleosome runs, all four tails were targeted.

**Table 4.3.**
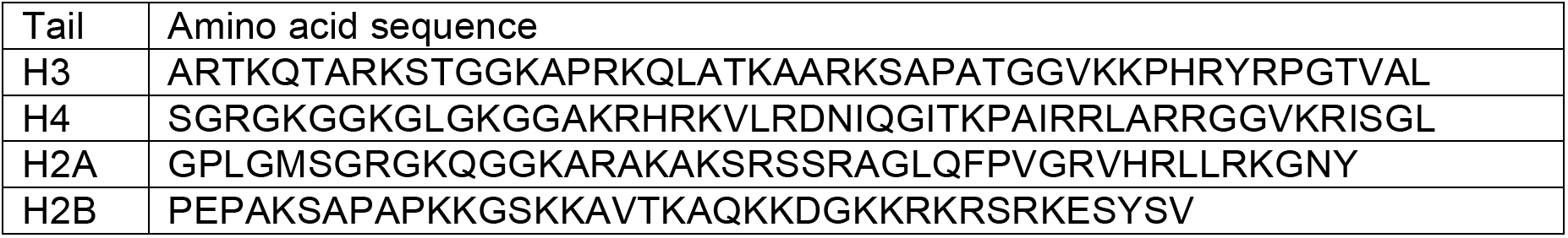
Amino acid sequences of tails analyzed in MD-HDX-MS/MS

**Table 4.4.** Mass, charge, and retention time information for tails analyzed in MD-HDX-MS/MS

| Tail | CS | RT (min) | Mass targeted for ETD |  |  |  |  |  |
| --- | --- | --- | --- | --- | --- | --- | --- | --- |
|  |  |  | ND | 10<sup>1</sup> s | 10<sup>2</sup> s | 10<sup>3</sup> s | 10<sup>4</sup> s | FD |
| H3 | +7 | 6-8.5 | 723.141 | 726.015 | 726.159 | 726.447 | 726.590 | 726.445 |
| H4 | +7 | 8.5-10 | 742.310 | 743.891 | 744.034 | 744.179 | 744.612 | 745.618 |
| H2A | +7 | 8.5-11 | 671.241 | 672.677 | 672.965 | 673.397 | 673.542 | 674.690 |
| H2B | +6 | 4.5-7 | 714.586 | 716.768 | 717.270 | 717.270 | 717.605 | 717.102 |

CS = charge state; ND = non-deuterated samples; FD = fully-deuterated samples

For non-deuterated samples, peptides were identified using pFind 3.0 for HCD data (peptide tolerance: 10 ppm, MS/MS tolerance: 0.4 Da)^40^ and Mascot for ETD data (peptide tolerance: 10 ppm, MS/MS tolerance: 0.04 Da)^41^ using a database containing the recombinant histone sequences. HDExaminer (Sierra Analytics, version 2.5) was used to analyze deuterium incorporation of peptides. For intact peptide analysis, the full sequence of the histone protein was imported into the software. The confident results of the pFind and Mascot searches were combined into a single CSV file and uploaded into the software as the “peptide pool.” Raw files were uploaded and analyzed using only the charge states identified by the pFind and Mascot searches.

HDExaminer was also used to analyze the MD-HDX-MS/MS tail peptide fragment ions. The sequence of the tail peptide was imported into the software (Table 4.3). Raw files were uploaded in top-down analysis mode, and the retention time was specified according to Table 4.3. Charge states +1 to the charge of the peptide were considered for analysis.

#### Scrambling analysis

To determine the degree of scrambling in the MD-HDX-MS/MS platform, we compared the deuterium content of the H2A tail fragment ions to the theoretical deuterium content of these ions with 100% scrambling. Theoretical values were calculated as previously described.^14^ A Kolmogorov-Smirnov two-sample test was used to test whether the difference in deuterium profiles of the experimental and theoretical fragment ions was statistically significant.

### Data access

All raw files are available on the Chorus Database (https://chorusproject.org/) (Project number 1471). All heat map data obtained from HDExaminer analysis is available in Supplemental Data 1. All calculated CV values are available in Supplemental Data 2.

## RESULTS

### Top-Down HDX-MS/MS

There are two major technical hurdles that must be addressed prior to conducting any HDX-MS/MS experiment: scrambling, whereby proton and deuterium atoms on the backbone can migrate, effectively randomizing signal, and back-exchange, where deuterium atoms on the protein can exchange for protons from desalting and infusion buffers.

Scrambling occurs when analyte ions reach high vibrational energy in the gas phase. Previous studies have indicated that using “gentle” MS conditions and fragmentation methods (such as ETD) can reduce scrambling to negligible levels. We monitored scrambling under various MS parameters using a probe peptide ‘P1’ (sequence: HHHHHHIIKIIK) as previously described and determined that scrambling levels are <10% under our given instrument parameters (isolation window: 10m/z, S lens RF: 60%, ETD reaction time: 70ms, resolution: 60,000, capillary temperature: 150°C) (Figure 2a).^8^ We found that the harshest conditions tested resulted in 63% scrambling (Isolation window: 2m/z, S lens RF: 70%, ETD reaction time: 150ms) (data not shown).

**Figure 2.**
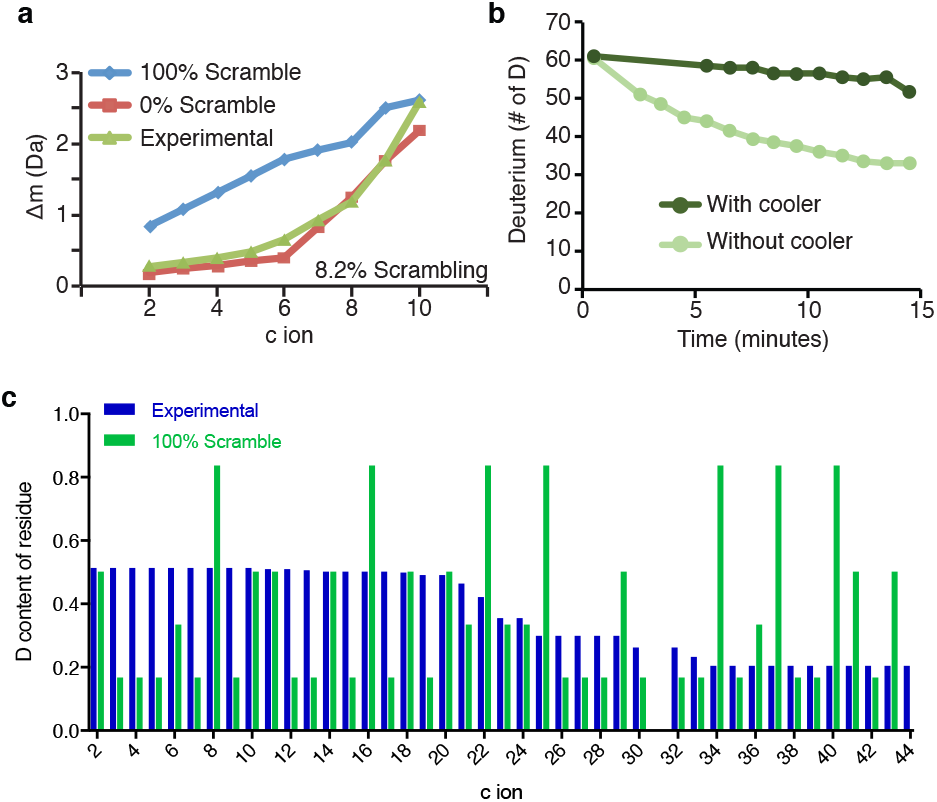
Scrambling and back-exchange can be minimized in the TD- and MD-HDX-MS/MS experiment. (a) Scrambling is minimial in the TD-HDX-MS/MS platform. The scrambling probe peptide ‘P1’ was used to monitor scrambling. The peptide was fragmented with ETD and the deuterium content of the resulting c ions was calculated. The non-deuterated centroid was subtracted from the deuterated centroid value to obtain the difference in mass (Δm) and plotted in green. The theoretical 100% scrambling (blue) and theoretical 0% scrambling (red) Δm values were calculated and plotted. (b) TD-HDX-MS/MS set-up with cooling apparatus reduces back-exchange. Fully-deuterated H2B sample was infused into the Thermo Orbitrap Fusion with and without the cooling apparatus. The deuterium content of the protein was monitored over a 15 minute infusion window. (c) Scrambling does not occur on the MD-HDX-MS/MS platform. The H2A tail peptide was targeted for fragmentation after 10s of deuterium exchange and the deuterium content of each fragment ion was calculated using HDExaminer (version 2.5). The theoretical deuterium content values assuming 100% scrambling was calculated for each fragment ion and compared to experimentally obtained values to determine if scrambling is occurring. A two-sample K-S test determined that the difference between the theoretical and experimental values are statistically significant (D = 0.291; D-crit = 0.144), indicating that scrambling is not occurring.

Our TD-HDX-MS/MS platform uses an Advion Triversa Nanomate to infuse samples. In this instrument, samples are picked up inside a plastic tip and brought to a silicon chip containing hundreds of nano-electrospray nozzles. These nozzles are next to the capillary, which is set to 170°C. To keep samples cool during infusion and minimize back-exchange, we constructed a home-made cooling apparatus as described by the Jørgensen group, where a cold stream of N_2_ gas is directed at the sample during infusion.^39^ To test this cooling apparatus, we infused fully-deuterated histone H2B and measured back-exchange as a shift in the centroid over a 15 minute infusion window. We found that back-exchange was reduced from 50% to less than 10% over the 15 minute window when using the cooling apparatus (Figure 2b).

Top-down HDX-MS/MS (TD-HDX-MS/MS) has not yet been conducted on a protein complex, and so we sought to optimize, expand, and apply TD-HDX-MS/MS methodology to accommodate histone H4 monomers and (H3/H4)_2_ heterotetramers. H4 monomers are likely to be mostly unstructured in the absence of an H3 binding partner, although no structural data is available. Conversely, although the crystal structure of the H3/H4 tetramer has not been solved, H4 within the tetramer likely contains the same three alpha helices that are observed in the nucleosome crystal structure based on previous HDX and EPR (electron paramagnetic resonance) spectroscopy data.^28,42,43^

Protection from deuterium exchange occurs when solvent does not have access to amide protons or when amide protons are occupied in hydrogen bonds, as they are in secondary structures. Therefore, slower rates of deuterium exchange can occur upon increased stability (or reduced flexibility) of protein regions including secondary structures, intra- or intermolecular contacts, compaction of the protein, or any combination of these. Therefore, we hypothesized that the unstructured H4 monomer would have faster rates of exchange compared to H4 in the context of a tetramer, which has substantial secondary structure and contact with H3.

To conduct this experiment, histones corresponding to the human sequences of H3.1 and H4 were expressed and purified in E. coli. Histone tetramers were reconstituted by salt dialysis. H4 monomers and (H3/H4)_2_ tetramers were incubated in D_2_O for varying timepoints, including 10^1^s, 10^2^s, 10^3^s, and 10^4^s, to allow for deuterium exchange for amide protons on the protein backbone. Experiments were performed in triplicate. After each time point, the exchange reaction was quenched and the sample was desalted at 4°C with homemade stage tip columns and subsequently flash-frozen. Samples were then thawed in ice water and infused using an Advion Triversa Nanomate into a Thermo Orbitrap Fusion instrument. The +15 charge state of histone H4 was fragmented using ETD, and deuterium content of the fragment ions were calculated using HDExaminer software (Sierra Analytics, version 2.5).

Our results demonstrate that the TD-HDX-MS/MS platform can easily distinguish between different types of secondary structures. We compared the exchange profiles of two H4 fragment ions in tetramer context, one located in the N-terminal tail domain (c15) and one located in the α-2 helix (z74) (Figure 3). We found that N-terminal tail fragment ion exhibited a small degree of protection from exchange, but was nearly fully deuterated by the last time point as is expected for protein regions lacking secondary structure. Conversely, we found that z74, which spans the α-3 helix and a portion of the α-2 helix exhibits a large degree of protection from exchange as expected. These results demonstrate the power of TD-HDX-MS/MS to monitor the stability of secondary structures.

**Figure 3.**
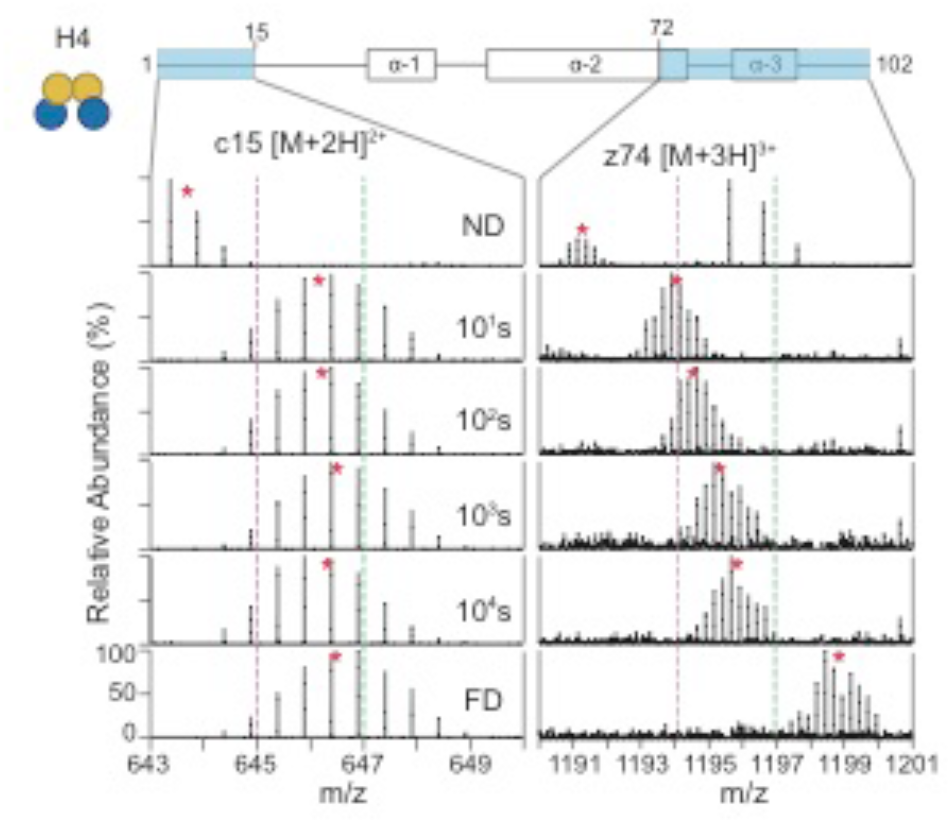
TD-HDX-MS/MS can distinguish different protein structures. The exchange profiles for c15 ([M+2H]^2+^) and z74 ([M+3H]^3+^) are shown for histone H4 in tetramer context across the HDX time course. Red stars denote the centroid of the isotopic distribution and the dotted purple and green lines act as guides to highlight the differences in m/z shifts. ND: non-deuterated; FD: fully deuterated.

The overall results of the TD-HDX-MS/MS experiment are displayed in Figure 4a and 4b. We were able to achieve highly resolved deuterium localization information for H4 in the monomer and tetramer constructs (28 or 36 site specific sites, respectively). Furthermore, we were able to obtain near site-specific resolution for the tail domains due to the identification of a large number of fragment ions in this region. We determined that the H4 tail domain has a slight degree of protection in the tetramer context, but is completely unstructured in monomeric H4 as full deuteration levels are reached by 10 seconds. We also obtained high coverage of the core region of H4 in both constructs, and we found that protected regions of H4 within the tetramer map to predicted secondary structures from the nucleosome crystal structure (Figure 4c). The structure shown contains Xenopus laevis histone sequences; however, the H4 sequence is identical to human and the H3 sequence is highly similar (99% identical).^25^ It is also important to note that the tail domains were not solved in the structure (electron density was too low) and the displayed tails were modeled in by the authors. Monomeric H4 was also found to contain regions of protection from deuterium exchange, particularly in the α-2 helix and α-3 helix (Figure 4a). These results demonstrate that TD-HDX-MS/MS can easily accommodate a histone complex to provide highly resolved exchange profiles, including the H4 tail domain.

**Figure 4.**
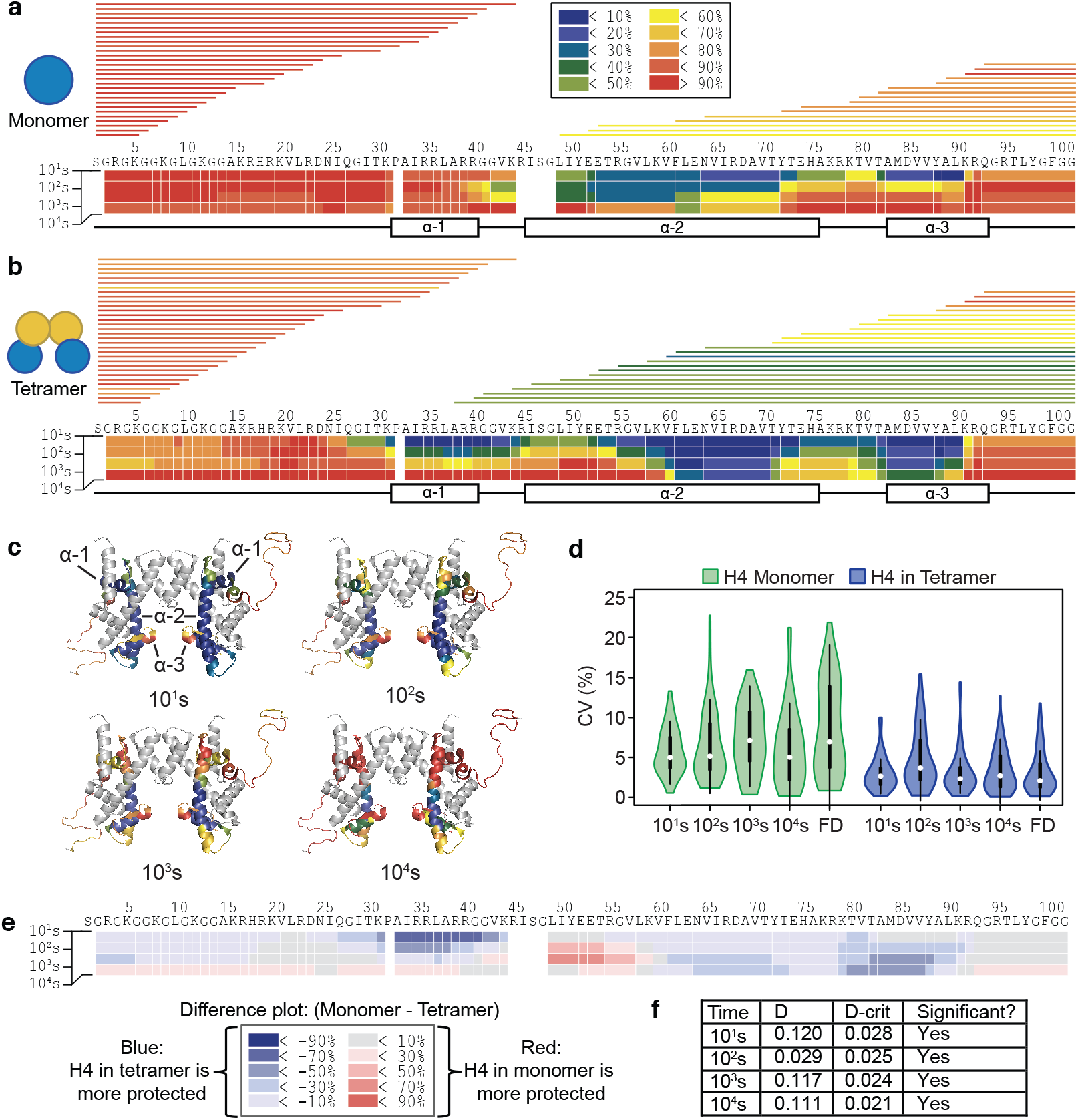
TD-HDX-MS/MS reveals that H4 undergoes different exchange profiles in the monomer compared to tetramer context. (a and b) Heat maps display the average % deuteration of each residue or group of residues according to the legend for H4 in (a) monomeric form or (b) within the tetramer from three experimental replicates. Each bar above the heat map indicates a fragment ion that was used for the analysis, and color of the bar indicates the % deuteration of that ion according to the legend. The secondary structure of H4 within the nucleosome is indicated below the heat map. (c) Regions of protection for H4 within the tetramer map to predicted secondary structures from the nucleosome crystal structure. Exchange profiles for H4 from the tetramer context are mapped to the crystal structure of the nucleosome (with DNA, H2A, and H2B hidden) (PDB: 1kx5). The color of the residues indicates the % deuteration for that residue as indicated by the legend. Histone H3 is colored in grey. (d) TD-HDX-MS/MS generates highly reproducible results. The CV values of the deuterium content of each fragment ion were calculated for the three experimental replicates and plotted. White circles show the medians; box limits indicate the 25th and 75th percentiles as determined by R software; whiskers extend 1.5 times the interquartile range from the 25th and 75th percentiles; polygons represent density estimates of data and extend to extreme values. Plot was generated using BoxPlotR.^44^ (e) H4 undergoes different exchange profiles as a monomer compared to tetramer. Deuterium % of the tetramer was subtracted from the monomer, and the results are shown according to the legend. Blue indicates that the tetramer is more protected and red indicates that the monomer is more protected from exchange. (f) Two-sample K-S test reveals that observed differences in H4 exchange profiles between monomer and tetramer are statistically significant. The maximum distance between the cumulative distribution of deuterium content (%) per residue of H4 in monomer and tetramer contexts was calculated (D) and compared to the calculated D-critical value (D-crit). Differences are significant if D is greater than D-crit.

It is important to note that H4 was found to have strikingly different exchange profiles in the monomer compared to tetramer (Figure 4e), consistent with a model where it co-folds with its partner histone, H3. A small region within the α-2 helix, spanning residues 52 to 57, experienced a larger degree of protection in the H4 monomer compared to tetramer; however, overall, H4 experienced a larger degree of protection in tetramer context compared to monomer. The regions undergoing the greatest degree of enhanced protection upon incorporation into the tetramer include the α-1 helix, the C-terminal half of the α-2 helix, and the α-3 helix. To determine whether the observed differences in exchange profiles of H4 in monomer compared to tetramer context are statistically significant, we performed a two-sample Kolmogorov-Smirnov (K-S) test. The K-S test is a non-parametric test that compares the cumulative distributions of two different data sets. To determine if the data sets are different, the K-S test calculates the largest difference between the two cumulative distributions (D) and compares this value to the largest difference tolerated under the null hypothesis (D-critical; D-crit), which states that the cumulative distributions are the same. Therefore if D is greater than D-critical, the null hypothesis is rejected, and the difference between the data sets is considered statistically significant. After performing the K-S test on our data, we found that the differences between the exchange profiles of H4 in monomer compared to tetramer contexts are statistically significant at each time point (Figure 4f).

We also assessed the reproducibility of deuterium content measurements on the TD-HDX-MS/MS platform. To this end, we analyzed the coefficient of variation (CV) of the deuterium content of each detected fragment ion between the three experimental replicates (Figure 4d). The results show that the CV values are very low, with the median CV value for the monomer and tetramer experiments being below 8% and 5%, respectively. These results indicate that the sample processing steps and the TD-HDX-MS/MS platform enables highly reproducible measurements of deuterium incorporation.

One inherent drawback of TD-HDX-MS/MS as conducted here is that omission of the RP-HPLC separation step limits the sample complexity that can be accommodated. If the sample is very complex, it becomes challenging to target a single protein for ETD due to spectral overlap with other species in the sample. If multiple species are selected together, the MS/MS spectrum will contain a lot of overlap from the many different fragment ions present, making analysis more difficult. Therefore, it will be very challenging to adapt this methodology to accommodate a more complex sample without the use of chromatography. Nonetheless, the results shown here demonstrate that TD-HDX-MS/MS is able to obtain robust and highly resolved deuterium localization information for intact proteins.

### Middle-down HDX-MS/MS

Bottom-up (BU) HDX-MS/MS employs digestion with pepsin and separation by RP-HPLC prior to analysis. As such, these platforms can handle more complex samples than TD- HDX-MS/MS, which does not typically employ separation techniques. Given that these platforms can handle more complex samples compared to TD-HDX-MS/MS, we sought to optimize the BU-HDX-MS/MS approach to accommodate (H3/H4)_2_ tetramers and nucleosomes. We targeted the histone tail domains for ETD fragmentation to obtain highly resolved exchange information. Given that these tail peptides are much longer than the standard peptide (ranging from 39 to 49 amino acids), we termed our platform middle-down (MD) HDX-MS/MS. This study represents the first heterogeneous protein complex and protein-DNA complex to be analyzed by MD-HDX-MS/MS. For comparison purposes, we analyzed these data sets without the ETD data (analogous to BU-HDX-MS) and with the ETD data (MD-HDX-MS/MS).

To this end, histones corresponding to the human sequences of H3.1, H4, and canonical H2A and H2B were expressed and purified in E. coli. A His-tag was engineered at the N-terminus of histone H2A to enable purification with a nickel column. A majority of the His-tag was cleaved off after purification, but a small 4-amino acid tag (sequence: GPLG) remains at the N-terminus. Histone tetramers or nucleosomes were reconstituted by salt dialysis. Nucleosomes were made with 197bp DNA sequence corresponding to the well-known 601 nucleosome positioning sequence.^45^ Tetramers and nucleosomes were incubated in D_2_O at 4°C for varying time points, including 10^1^s, 10^2^s, 10^3^s, and 10^4^s, to allow for deuterium exchange of amide protons on the protein backbone. Experiments were performed in at least triplicate. Each tail was identified in the non-deuterated control and was targeted for ETD fragmentation in deuterated sample runs. Results were analyzed using HDExaminer software (Sierra Analytics, version 2.5).

Given that the MS set-up for MD-HDX-MS/MS is different than that for TD-HDX-MS/MS, we first sought to ensure that scrambling was not occurring on our platform. To this end, we analyzed the H2A tail peptide, which has the highest charge density of the peptides analyzed and is therefore the most likely to scramble. We calculated the theoretical deuterium content of each fragment ion under 100% scrambling conditions at the 10s time point based on how many labile hydrogen atoms are present as has been done by the Williams group.^14^ We plotted these values along with the experimentally obtained deuterium content values (Figure 2c). To determine whether the experimental and theoretical values are different, we performed a two-sample Kolmogorov-Smirnov (K-S) test and found that the difference between the theoretical and experimental deuterium content values are statistically significant (D = 0.291, D-crit = 0.144), indicating that scrambling is not occurring in our MD-HDX-MS/MS platform.

The results of the MD-HDX-MS/MS analysis are shown in Figure 5. Notably, the peptides spanning the tail domains are very long for each histone and so highly resolved information for this region would not be possible without the use of ETD (Supplemental Figure 1). However, we were able to successfully target a peptide spanning the histone tail domain for each histone and obtain highly resolved deuterium localization information (Figure 5). The H3 and H4 tail peptides had higher fragment ion coverage than H2A and H2B, likely due to their higher abundance.

**Figure 5.**
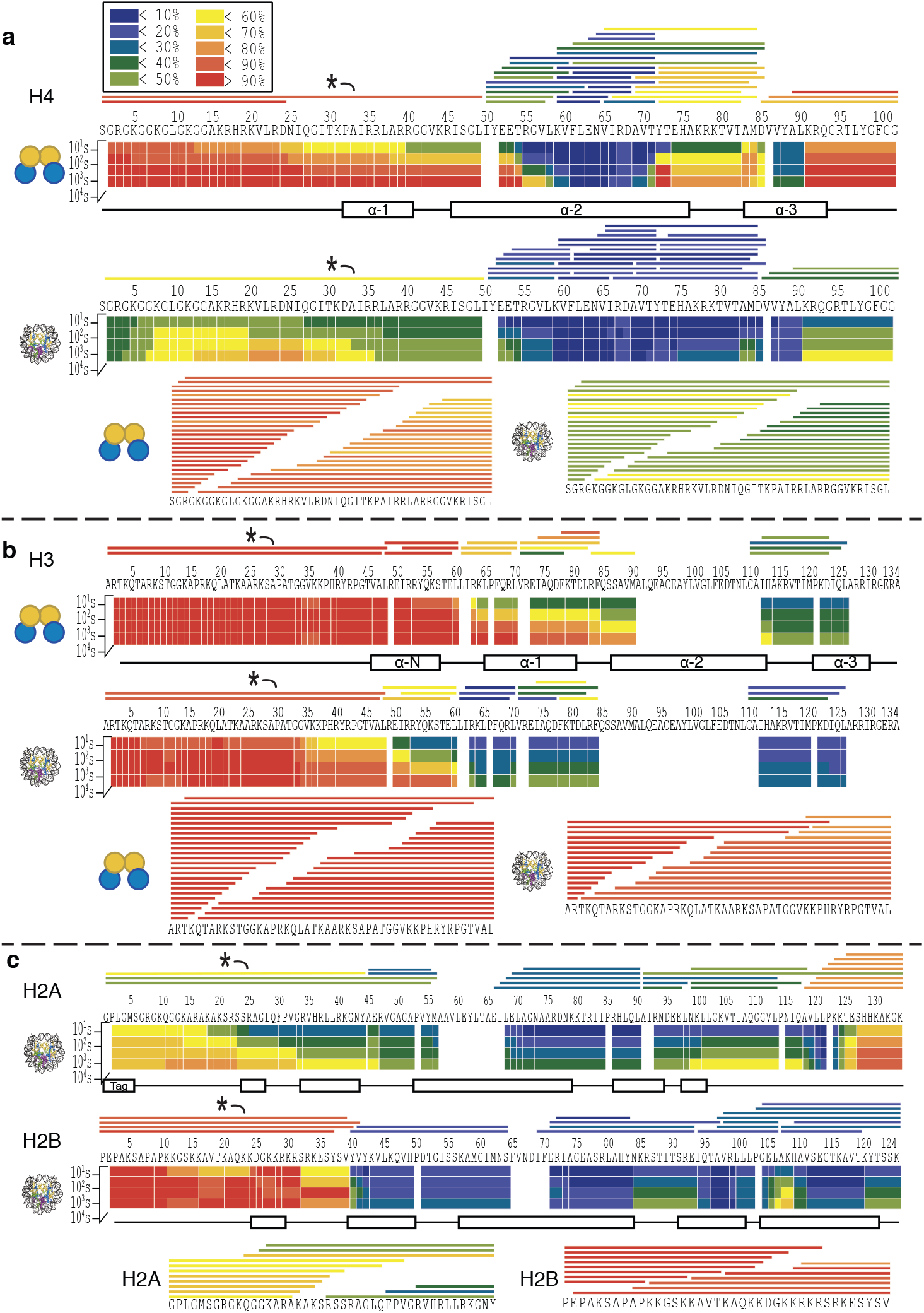
MD-HDX-MS/MS platform enables near-site specific resolution of deuterium localization in histone tail domains. Exchange profiles are given for H4 (a), H3 (b), H2A and H2B (c) in tetramer (when relevant) and nucleosome contexts as demonstrated by the icon on the left. The heat maps display the average % deuteration of each residue or group of residues according to the legend. The bars above the heat maps indicate peptides that were used in the analysis. The peptide labeled with a star (*) was selected for ETD fragmentation, and the fragment ions used in the analysis are shown below the heat maps. The color of each bar indicates the average % deuteration according to the legend.

The regions of protection observed in the core domain of each histone align well with the secondary structures found in the crystal structure (Figure 5 and Supplemental Figure 2). Histone H4 demonstrates a large degree of protection in the α-helices; however, the α-2 and α-3 helices experience a greater degree of protection than the α-1 helix in both tetramer and nucleosome contexts (Figure 5a). The α-N helix of H3 is completely unstructured in tetramer context as indicated by extremely rapid deuterium exchange, but is protected in the nucleosome context, indicating that this helix may be stabilized by interaction with DNA and/or the presence of H2A/H2B dimers within the nucleosome (Figure 5b). There is a large degree of protection in the alpha helices of H2A and H2B in nucleosomal context, and this protection extends into the linker regions between helices (Figure 5c). The H2B tail appears to be mostly unstructured, as it is nearly fully deuterated by 10 seconds (deuterium content ranges from 82% to 96% in the tail throughout the time course). Conversely, the H2A tail experiences a large degree of protection form exchange, experiencing a maximum of 51% deuterium incorporation at 10 seconds and reaching a maximum of 69% deuterium incorporation by 10^4^ seconds.

Overall, H3 and H4 in nucleosomal context are more protected from exchange compared to the tetramer context, indicating that incorporation into the nucleosome stabilizes H3 and H4 as has been observed previously (Figure 6a-c).^31,42^ Enhanced protection of H4 occurs globally, with virtually the entire protein undergoing an increase in protection in the nucleosome compared to tetramer (Figure 6b). The H4 tail undergoes a dramatic increase in protection from exchange in the nucleosome compared to tetramer, reaching differences in deuterium content of up to 64% between the two constructs (Supplemental Figure 3). For H3, the core region of the protein also experiences an increase in protection in the nucleosome compared to tetramer (Figure 6c). The H3 tail exhibits some increased protection within the nucleosome, mainly corresponding to the α-N helix, which reaches up to 44% greater protection in the nucleosome compared to the tetramer. The region immediately adjacent to the α-N helix on the N-terminal side also experiences an increase in protection from exchange, spanning approximately 10 amino acids. Nearly the entire tail undergoes slight protection (around 10% lower deuterium content) in the 10 second time point, indicating that incorporation into the nucleosome confers a small degree of protection from exchange for the H3 tail relative to the tetramer, however the most dramatic difference is located in the α-N helix and region immediately adjacent. Together, these results show that incorporation into the nucleosome confers increased structural rigidity to the H3 and H4 core as well as the tails, albeit to different degrees.

**Figure 6.**
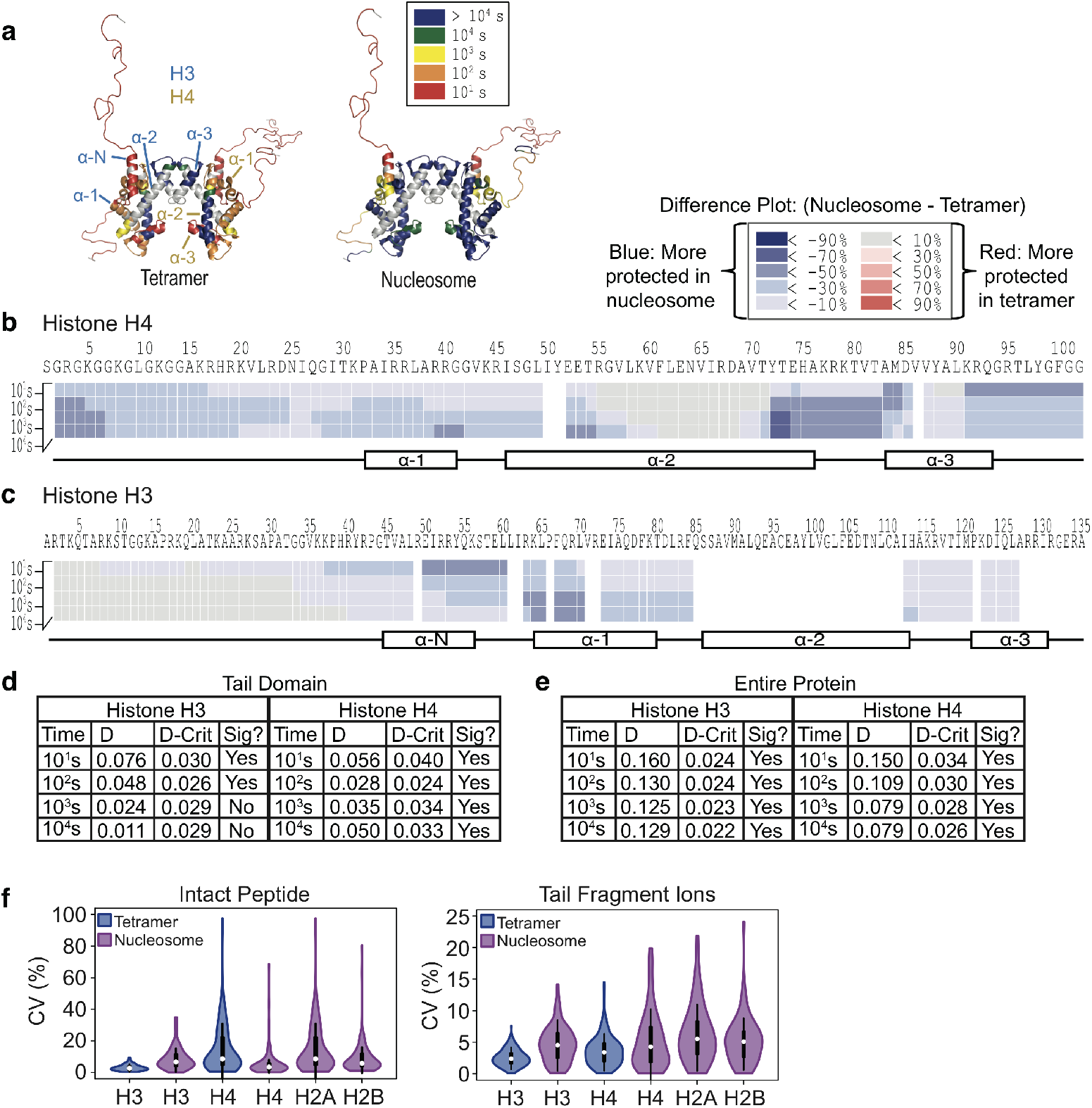
H3 and H4 exhibit increased protection from exchange upon incorporation into the nucleosome. (a) Increased protection of H3 and H4 in the nucleosome compared to tetramer occurs globally. The exchange profiles for H3 and H4 from the tetramer and nucleosome contexts as measured by MD-HDX-MS/MS are mapped to the crystal structure of the nucleosome (with DNA, H2A, and H2B hidden for easier viewing) (PDB: 1kx5). The color of each residue indicates the time point at which that residue was greater than 50% deuterated. Gray color indicates that no information was available due to lack of sequence coverage for that region. Secondary structures of H3 (blue) and H4 (yellow) are indicated. (b and c) Difference plots for H4 (b) and H3 (c). Deuterium % of the tetramer was subtracted from the nucleosome, and the results are shown according to the legend. Blue indicates that the nucleosome is more protected and red indicates that the tetramer is more protected from exchange. Secondary structures are mapped underneath the difference plot. (d and e) two-sample K-S test indicates that differences in exchange for H3 and H4 in tetramer compared to nucleosomal context are statistically significant for (d) the tail domains and (e) the entire protein. The maximum distance between the cumulative distribution of deuterium content (%) per residue of H4 in tetramer and nucleosome contexts (D) was calculated and compared to the calculated D-critical value (D-crit). Differences are considered statistically significant if D is greater than D-crit. (f) MD-HDX-MS/MS enables highly reproducible deuterium content measurements. The CV values of the deuterium content of intact peptides and tail peptide fragment ions were calculated across all time points and plotted. White circles show the medians; box limits indicate the 25th and 75th percentiles as determined by R software; whiskers extend 1.5 times the interquartile range from the 25th and 75th percentiles; polygons represent density estimates of data and extend to extreme values. Plot was generated using BoxPlotR.^44^

To determine whether the observed differences in exchange profiles for the H3 and H4 tails in the tetramer compared to nucleosome contexts are statistically significant, we performed a two-sample K-S test for each time point (Figure 6d-e). The results indicate that the H4 tail has a significantly different exchange profile in tetramer compared to nucleosomal contexts across all time points. However, only the first two time points of the H3 exchange profile have a statistically significant difference, likely because the tail is nearly fully deuterated in both constructs by the last two time points (Figure 6d). We also performed a K-S two-sample test comparing the exchange profiles of the entire H3 and H4 protein, and the results indicate that, overall, the differences in exchange profiles of H3 and H4 in tetramer compared to nucleosome contexts are statistically significant across all time points (Figure 6e).

We also assessed the reproducibility of deuterium content measurements for the tail fragment ions as well as the intact peptides between the experimental replicates (3 to 5 replicates for each time point). To this end, we calculated the CV of the deuterium content measurement for each detected fragment ion or peptide across all time points (Figure 6f). We were able to achieve highly reproducible deuterium content measurements, with the median CV value of the fragment ions being below 7% for each protein and the median value of the intact peptides being below 10% for each protein, indicating that the MD-HDX-MS/MS platform generates highly reproducible results.

### Evaluation and comparison of BU-HDX-MS, MD-HDX-MS/MS, and TD-HDX-MS/MS platforms

We next sought to compare the results obtained on the MD- and TD-HDX-MS/MS platforms to determine how well the results agree. To this end, we compared the exchange profiles of H4 in tetramer context as this was the only protein analyzed on both platforms (Figure 7a). Overall, the exchange profiles are highly similar, with regions of secondary structure exhibiting increased protection relative to other regions of the protein. We performed a Pearson correlation analysis for each time point based on the deuterium content of each amino acid (Figure 7b). We found that each time point demonstrates a high degree of similarity between TD and MD platforms, with the following correlation coefficients: 10s: 0.837; 100s: 0.847; 1,000s: 0.842; 10,000s: 0.883. Each comparison was found to have statistically significant correlation values, with p-values less than 6 x 10^-23^. We also performed a Pearson correlation analysis for all time points together and obtained an overall correlation coefficient of 0.858 (p-value: 1.71 x 10^-29^), indicating that the TD- and MD-HDX-MS/MS platforms yielded highly similar results.

**Figure 7.**
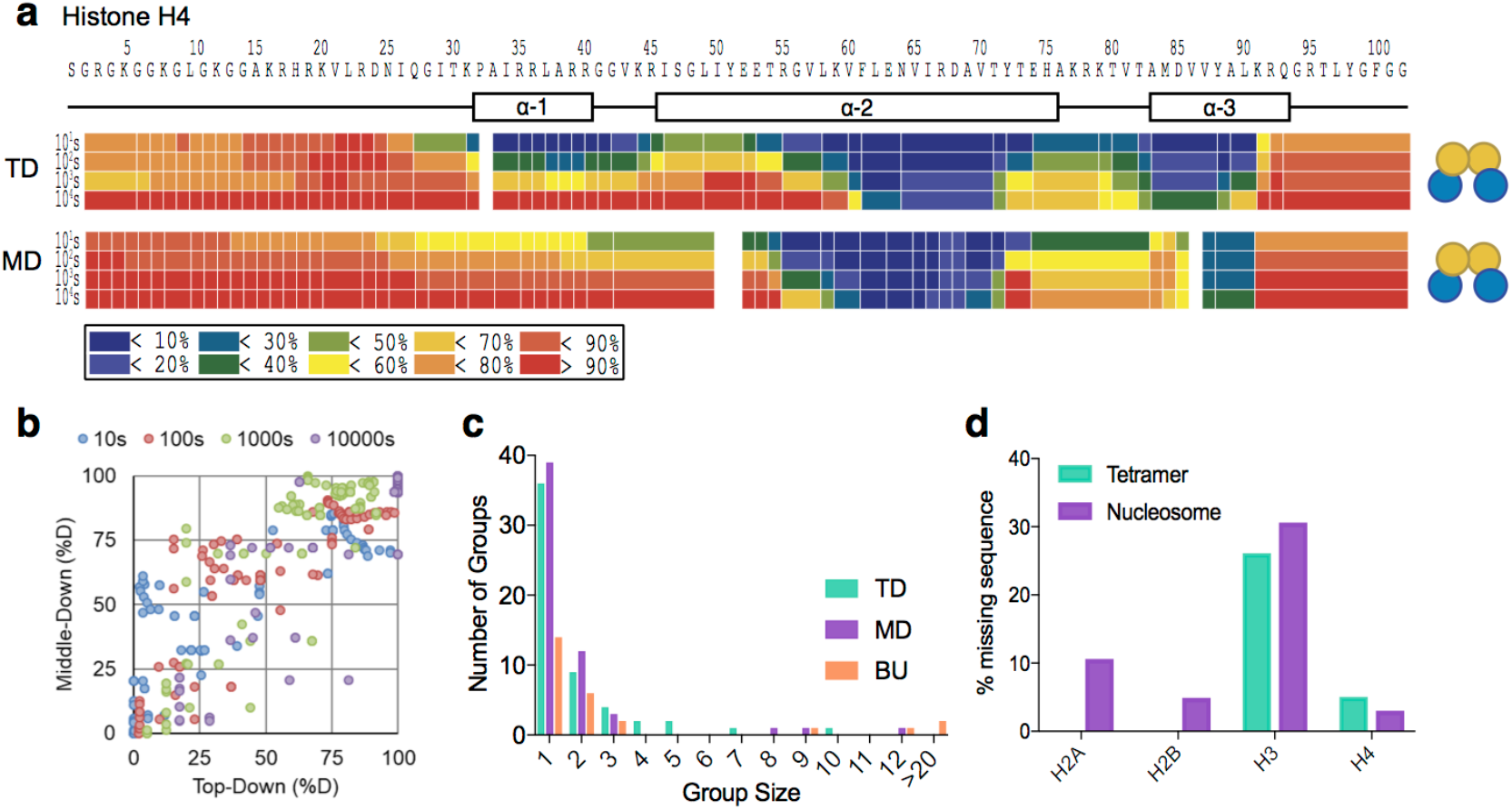
A comparison of the BU-HDX-MS, MD-HDX-MS/MS, and TD-HDX-MS/MS platforms. (a) TD- and MD-HDX-MS/MS platforms provide comparable deuterium content measurements for histone H4 in tetramer context. Heat maps indicate the % deuteration of that residue according to the legend below the figure. The secondary structure is displayed above the heat maps. (b) Correlation plot for deuterium content of H4 in tetramer context for TD- and MD-HDX-MS/MS. Each point on the plot is color coded by time point and represents the deuterium content of a single residue. A Pearson correlation was performed for each time point with the following coefficients: 10s: 0.837; 100s: 0.847; 1,000s: 0.842; 10,000s: 0.882; all time points together: 0.858. The p-values were less than 6 x 10^-27^ for each correlation. (c) TD- and MD-HDX-MS/MS afford much greater resolution of deuterium content localization for histone H4 in tetramer context compared to BU-HDX-MS. Group size indicates the number of residues that were analyzed together (i.e. 1 is site-specific information, 2 indicates that two amino acids were analyzed together, etc.). (d) MD-HDX-MS/MS results in sequence coverage gaps. The percent of amino acid sequence in which no information was obtained due to incomplete coverage of identified peptides is shown for each histone in nucleosome context and tetramer context (H3 and H4 only).

One of the greatest advantages of TD-HDX-MS/MS is that it has the potential to achieve site-resolved information across the entire protein sequence, depending on the quality of fragmentation. In MD-HDX-MS/MS, on the other hand, only certain peptides will be amenable to ETD fragmentation. In this case, only the tail peptides had a high enough charge state and efficient fragmentation to be analyzed with ETD. Therefore, the other regions of the protein can only be studied at the intact level and will be dependent upon the presence of many overlapping peptides to get more site-resolved information, which is not always possible depending on the protein sequence. We analyzed the differences in resolution between the MD- and TD-HDX-MS/MS platforms for H4 in tetramer form as well as the MD-HDX-MS/MS data without including the ETD data, which is analogous to a BU-HDX-MS experiment (Figure 7c). To compare resolution between platforms, we determined how many residues had to be grouped together for analysis. The results demonstrate that the MD-HDX-MS/MS platform had a slightly higher number of singly resolved sites compared to TD-HDX-MS/MS (39 and 36, respectively); however, MD-HDX-MS/MS had larger groups of residues that had to be analyzed together (1 group of 8, 9, and 12 amino acids) compared to TD-HDX-MS/MS (1 group of 5, 7, and 10). As expected, BU-HDX-MS resulted in the worst level of resolution, with only 14 singly-resolved sites and more large groups (two of which are greater than 20 amino acids), highlighting the power of ETD-based HDX methods to improve the resolution of deuterium localization information compared to traditional BU-HDX-MS approaches.

Despite the fact that the resolution was highly similar between MD-HDX-MS/MS and TD-HDX-MS/MS for histone H4, the TD platform has the advantage of guaranteeing full sequence coverage. The H4 protein had near complete coverage in MD-HDX-MS/MS; however, the other histone proteins had a greater amount of gaps in sequence coverage, up to 30% for H3 (Figure 7d). Therefore, while the resolution may be similar for H4, TD-HDX-MS/MS may be desirable for proteins in which MD-HDX-MS/MS does not yield full coverage.

## Discussion

The results described here demonstrate the power of ETD-based HDX methods to obtain deuterium localization information in great detail. We were able to analyze, for the first time, a protein complex with TD-HDX-MS/MS and a heterogeneous protein complex and protein/DNA complex with MD-HDX-MS/MS. We were able to increase the number of site-specific deuterium localization from 14 sites in BU-HDX-MS to 36 and 39 sites in TD-HDX-MS/MS and MD-HDX-MS/MS, respectively for histone H4 in tetramer context. The deuterium content measurements on each platform were highly reproducible, with the median coefficient of variation (CV) values of fragment ions and intact peptides being below 10% for both platforms. Together, these results show that MD- and TD-HDX-MS/MS are capable of rigorous, reproducible, and highly resolved deuterium content measurements. Furthermore, these HDX-MS/MS platforms can be tailored to target specific proteins or peptides of interest for ETD and can therefore be easily applied to any sample, demonstrating the wide utility and versatility of these methods.

Here, we develop and use these ETD-based HDX platforms to provide the first detailed view of histone tail dynamics in different contexts. We first analyzed histone H4 in monomer and (H3/H4)_2_ heterotetramer contexts with TD-HDX-MS/MS and were able to obtain highly resolved information across the entire sequence of H4. We next analyzed (H3/H4)_2_ heterotetramers and intact nucleosomes on our MD-HDX-MS/MS platform. We targeted each tail for fragmentation, enabling highly resolved information to be obtained for each histone tail. The remainder of the protein was analyzed at the intact peptide level. However, because many overlapping peptides were obtained in the digestion, we were able to match the resolution afforded by TD-HDX-MS/MS for histone H4 in tetramer context (Figure 7c).

Our TD- and MD-HDX-MS/MS results are corroborated by previous crystallography and BU-HDX-MS studies, indicating that this method is highly accurate and reliable. Observed areas of protection for each histone overlap nearly exactly with the secondary structures observed in the crystal structure of the nucleosome (Figure 4c and Supplemental Figure 2).^28^ It is important to note that no structural information is available for the H4 monomer; however, we found that the H4 monomer also contains some areas of protection, mainly in the α-2 helix, indicating that the monomer may have some secondary structure or form protein complexes, although it cannot be ruled out that some aggregation occurred during sample processing. The tail domains were not fully solved in the crystal structure due to low electron density in those regions, precluding a comparison of histone tail domains.

Previous BU-HDX-MS analyses of histone tetramers and nucleosomes also corroborate our findings.^31,42^ H3 and H4 core domains undergo a massive increase in stability upon incorporation into the nucleosome compared to tetramer (> 3 orders of magnitude), which we also observed in our MD-HDX-MS/MS results. Furthermore, we found that the α-2 and α-3 helices of H4 exhibit a larger degree of protection than the α-1 helix within the tetramer, as was also observed in the earlier HDX data.^31,42^ We also found that the α-N helix of H3 is highly unstructured in the tetramer as this region experiences nearly 100% deuterium incorporation by 10 s. However, our data show that the α-N helix undergoes an increase in protection from exchange within the nucleosome, suggesting that this helix is stabilized upon incorporation into the nucleosome, a finding which was also observed in the previous BU-HDX-MS studies.^31,43^ Overall, our results match previous crystallography and BU-HDX-MS data for the core domains of histones, demonstrating that the TD- and MD-HDX-MS/MS platforms provide reliable deuterium content measurements and insights into protein structure and dynamics.

Although these previous studies lend significant insight into histone core structure and stability, they were unable to provide detailed analysis of histone tail domains. The tail domains have not been fully solved in crystallography studies due to low electron density measurements, likely due to the flexible nature of the histone tails. Furthermore, BU-HDX-MS has been unable to provide a detailed analysis of histone tails, either because they were not detected or because the peptides corresponding to the tail domain were too long to provide resolved information likely due to the high content of lysine and arginine residues present in the tail domains.

Here, our TD- and MD-HDX-MS/MS platforms are able to provide the first detailed analysis of histone tail dynamics in solution in nucleosome and sub-nucleosomal contexts. We found that the H3 tail is completely disordered in tetramer context, reaching near 100% deuteration by 10 seconds, including the region corresponding to the first half of the α-N helix. However, we found that the H3 tail undergoes an increase in protection in the nucleosomal context compared to tetramer that was localized to the α-N helix and a 10 amino acid stretch immediately adjacent to the helix on the N-terminal side (Figure 5b). This 10 amino acid section aligns with the portion of the tail that exits the nucleosomal core between the two DNA superhelical gyres close to the dyad axis, indicating that this protection may be due to interaction with the DNA (Figure 8a). Indeed, crosslinking studies indicate that the H3 tail can interact with DNA approximately 35-40 bp from the dyad^46^, precisely where the tail is mapped in the crystal structure.^28^ Note that the tail domains were not solved in the crystal structure as the electron density was too low but rather modeled in afterwards.

**Figure 8.**
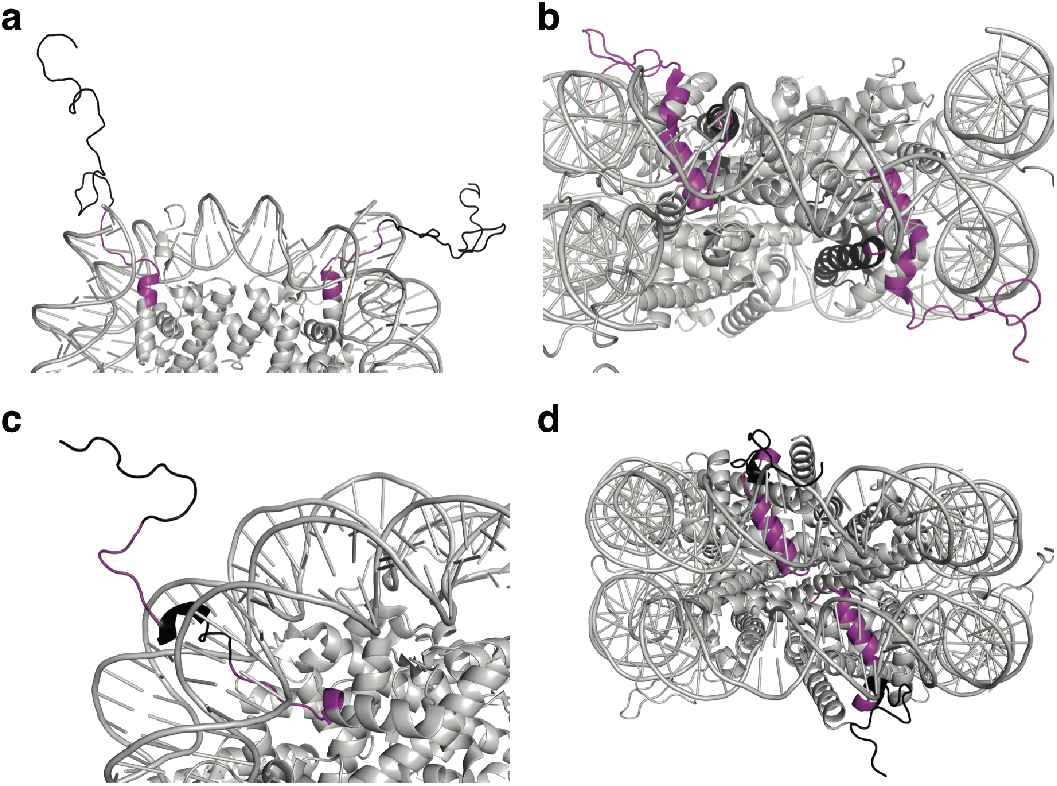
The histone tail domains are protected from exchange in nucleosomal context. Each tail peptide analyzed in MD-HDX-MS/MS is colored black, with the area of greatest protection at 10s are colored in purple for H3 (A), H4 (B), H2B (C), and H2A (D). For H3, amino acids 35-48 are colored, which exhibit approximately 60% deuterium incorporation. For H4, residues 27-49 are colored, which exhibit approximately 40% deuterium incorporation. For H2B, residues 16-23 and 32-39 are colored, which exhibit approximately 60% deuterium incorporation. For H2A, residues 25-44 are colored, which exhibit less than 30% deuterium incorporation. PDB: 1kx5. Note that the tail domains in this structure were modeled in after solving the structure because no electron density could be assigned to them.

The H4 tail undergoes a dramatic increase in protection upon incorporation into the nucleosome relative to the tetramer. Within the tetramer, the H4 tail exhibits protection corresponding to the α-1 helix of the sequence, although this protection extends into the tail domain as well, albeit to a lesser degree. Within the nucleosome, however, the tail exhibits a large degree of protection, only reaching between 40-80% deuteration levels by the last time point, indicating that the nucleosome context reduces the conformational flexibility of the tail (Figure 5a). The H4 tail protrudes from the nucleosomal surface from under the DNA superhelical gyre, placing it in a position where it could potentially interact with the nucleosomal DNA (Figure 8b). Indeed, crosslinking studies have demonstrated that the H4 tail can interact with DNA approximately 35-40 bp from the dyad.^47^ However, previous studies have also shown that residues 16-24 of the histone tail can bind to a region on the H2A and H2B surface called the acidic patch, named for the 8 acidic residues located at this surface.^28,48^ Most of these studies have shown the H4 tail binding to the acidic patch of an adjacent nucleosome, an interaction which has been shown to be critical in chromatin compaction and folding into higher order structures.^28,49^ However, Kan and colleagues also determined, using crosslinking strategies on mono-nucleosomes, that an H4 tail can bind the acidic patch within the same nucleosome.^50^ A third possibility is that the H4 tail may contain secondary structure in nucleosomal context, such as an alpha helix. Further experiments are needed to determine which of these interactions is occurring in solution.

The H2A and H2B tail peptides were studied in nucleosomal context only. The H2B peptide exhibits some protection corresponding to the area containing the first α helix as expected but also contains an additional area of slight protection in the middle of the tail domain (Figure 5d). This result indicates that there may be some secondary structure present or that this region interacts with the nucleosome. Indeed, the secondary structure prediction algorithm, PSIPRED, demonstrated that this region has a propensity for alpha-helix formation.

Furthermore, the tails within the crystal structure do harbor a small α-helix in the tail domain, nearby the observed area of protection, although they do not overlap. It is important to note that the tail was not solved in the crystal structure, but rather modeled in afterwards (Figure 8c). Previous bottom-up HDX data from the Luger lab demonstrated that H2A and H2B in dimer context are nearly completely exchanged by 10s, indicating that these proteins likely undergo a massive increase in structural rigidity upon incorporation into the nucleosome.^51^

The H2A tail peptide, on the other hand, experiences a large degree of protection from deuterium exchange in nucleosomal context, reaching a maximum of 80% deuteration levels by the last time point (10^4^ seconds). This protection spans the entire length of the H2A tail as well as the first alpha-helix of the protein. The greatest protection in the tail domain is localized to the region immediately adjacent to the alpha-helix on the N-terminal side (Figure 5c). The H2A tail exits the nucleosomal surface underneath the DNA superhelical gyre in a position where the tail could potentially interact with the nucleosomal DNA (Figure 8d). This result is corroborated by the Hayes group who used crosslinking to show that residue A12 and G2 of the H2A tail can crosslink to the nucleosomal DNA 40 bp and 35-45 bp from the nucleosomal dyad, respectively.^52^

We also compared the performances of our BU-HDX-MS, MD-HDX-MS/MS, and TD-HDX-MS/MS for histone analysis. To this end, we compared the deuterium localization results of histone H4 in tetramer context, as this is the only protein that was analyzed across all three platforms (Figure 7). The results demonstrate that all three platforms generate highly similar results, although MD- and TD-HDX-MS/MS enable far superior resolution compared to BU-HDX-MS. For histone H4, TD- and MD-HDX-MS/MS yielded similar levels of resolution. This result is likely due to the fact that the pepsin digest for H4 yielded many overlapping peptides, enabling more site-specific deuterium localization to be obtained. However, full coverage for the other histone proteins was not obtained in the MD-HDX-MS/MS platform, preventing analysis of these regions using this platform. TD-HDX-MS/MS ensures full coverage of the protein of interest and is therefore not a limitation in this approach.

Given that these methods returned highly similar results, determining the optimal platform for a protein of interest will depend upon the exact experimental conditions because each of the platforms has different limitations and strengths. TD-HDX-MS/MS ensures full coverage of the protein, requires dramatically less instrument time, has fewer sample processing steps, and can achieve higher resolution than MD-HDX-MS/MS. However, because the digestion and separation steps are omitted in this platform, samples must be relatively simple to avoid spectral overlap. Furthermore, the protein of interest must be ETD-amenable or the resolution obtained will be poor. Given that ETD operates with an electron transfer mechanism, charge-dense proteins will be more amenable to this technique. MD-HDX-MS/MS, on the other hand, can accommodate more complex samples and requires approximately 5-to 10-fold less protein. Additionally, the protein does not have to be ETD-amenable because any charge-poor regions of the protein can be analyzed at the intact peptide level.

Overall, these results demonstrate that MD-HDX-MS/MS and TD-HDX-MS/MS enable precise and highly reproducible deuterium localization of proteins in complex samples. This study represents the first heterogeneous protein complex and protein/DNA complex to be analyzed with ETD-based HDX-MS methodology, demonstrating the versatility and power of these methods for extremely detailed structural and dynamic analysis of protein molecules.

## Acknowledgements

We gratefully acknowledge funding from NIH grant R01-GM105654 (B.E.B).

## References

(1) Althaus, E.; Canzar, S.; Ehrler, C.; Emmett, M. R.; Karrenbauer, A.; Marshall, A. G.; Meyer-Bäse, A.; Tipton, J. D.; Zhang, H.-M. Computing H/D-Exchange Rates of Single Residues from Data of Proteolytic Fragments. BMC Bioinformatics 2010, 11, 424.

(2) Mayne, L.; Kan, Z.-Y.; Chetty, P. S.; Ricciuti, A.; Walters, B. T.; Englander, S. W. Many Overlapping Peptides for Protein Hydrogen Exchange Experiments by the Fragment Separation-Mass Spectrometry Method. J. Am. Soc. Mass Spectrom. 2011, 22 (11), 1898.

(3) Gessner, C.; Steinchen, W.; Bédard, S.; J. Skinner, J.; Woods, V. L.; Walsh, T. J.; Bange, G.; Pantazatos, D. P. Computational Method Allowing Hydrogen-Deuterium Exchange Mass Spectrometry at Single Amide Resolution. Sci. Rep. 2017, 7.

(4) Kan, Z.-Y.; Walters, B. T.; Mayne, L.; Englander, S. W. Protein Hydrogen Exchange at Residue Resolution by Proteolytic Fragmentation Mass Spectrometry Analysis. Proc. Natl. Acad. Sci. 2013, 110 (41), 16438–16443.

(5) Rand, K. D.; Zehl, M.; Jensen, O. N.; Jørgensen, T. J. D. Protein Hydrogen Exchange Measured at Single-Residue Resolution by Electron Transfer Dissociation Mass Spectrometry. Anal. Chem. 2009, 81 (14), 5577–5584.

(6) Abzalimov, R. R.; Kaplan, D. A.; Easterling, M. L.; Kaltashov, I. A. Protein Conformations Can Be Probed in Top-Down HDX MS Experiments Utilizing Electron Transfer Dissociation of Protein Ions Without Hydrogen Scrambling. J. Am. Soc. Mass Spectrom. 2009, 20 (8), 1514–1517.

(7) Hamuro, Y.; Tomasso, J. C.; Coales, S. J. A Simple Test To Detect Hydrogen/Deuterium Scrambling during Gas-Phase Peptide Fragmentation. Anal. Chem. 2008, 80 (17), 6785–6790.

(8) Rand, K. D.; Jørgensen, T. J. D. Development of a Peptide Probe for the Occurrence of Hydrogen (1H/2H) Scrambling upon Gas-Phase Fragmentation. Anal. Chem. 2007, 79 (22), 8686–8693.

(9) Zehl, M.; Rand, K. D.; Jensen, O. N.; Jørgensen, T. J. D. Electron Transfer Dissociation Facilitates the Measurement of Deuterium Incorporation into Selectively Labeled Peptides with Single Residue Resolution. J. Am. Chem. Soc. 2008, 130 (51), 17453–17459.

(10) Huang, R. Y.-C.; Garai, K.; Frieden, C.; Gross, M. L. Hydrogen/Deuterium Exchange and Electron-Transfer Dissociation Mass Spectrometry Determine the Interface and Dynamics of Apolipoprotein E Oligomerization. Biochemistry (Mosc.) 2011, 50 (43), 9273–9282.

(11) Landgraf, R. R.; Chalmers, M. J.; Griffin, P. R. Automated Hydrogen/Deuterium Exchange Electron Transfer Dissociation High Resolution Mass Spectrometry Measured at Single-Amide Resolution. J. Am. Soc. Mass Spectrom. 2012, 23 (2), 301–309.

(12) Abzalimov, R. R.; Bobst, C. E.; Kaltashov, I. A. A New Approach to Measuring Protein Backbone Protection with High Spatial Resolution Using H/D Exchange and Electron Capture Dissociation. Anal. Chem. 2013, 85 (19), 9173–9180.

(13) Pan, J.; Zhang, S.; Borchers, C. H. Comparative Higher-Order Structure Analysis of Antibody Biosimilars Using Combined Bottom-up and Top-down Hydrogen-Deuterium Exchange Mass Spectrometry. Biochim. Biophys. Acta BBA - Proteins Proteomics 2016, 1864 (12), 1801–1808.

(14) Masson, G. R.; Maslen, S. L.; Williams, R. L. Analysis of Phosphoinositide 3-Kinase Inhibitors by Bottom-up Electron-Transfer Dissociation Hydrogen/Deuterium Exchange Mass Spectrometry. Biochem. J. 2017, 474 (11), 1867–1877.

(15) Xiao, H.; Kaltashov, I. A. Transient Structural Disorder as a Facilitator of Protein-Ligand Binding: Native H/D Exchange—Mass Spectrometry Study of Cellular Retinoic Acid Binding Protein I. J. Am. Soc. Mass Spectrom. 2005, 16 (6), 869–879.

(16) Hoerner, J. K.; Xiao, H.; Kaltashov, I. A. Structural and Dynamic Characteristics of a Partially Folded State of Ubiquitin Revealed by Hydrogen Exchange Mass Spectrometry. Biochemistry (Mosc.) 2005, 44 (33), 11286–11294.

(17) Pan, J.; Han, J.; Borchers, C. H.; Konermann, L. Electron Capture Dissociation of Electrosprayed Protein Ions for Spatially Resolved Hydrogen Exchange Measurements. J. Am. Chem. Soc. 2008, 130 (35), 11574–11575.

(18) Pan, J.; Han, J.; Borchers, C. H.; Konermann, L. Hydrogen/Deuterium Exchange Mass Spectrometry with Top-Down Electron Capture Dissociation for Characterizing Structural Transitions of a 17 KDa Protein. J. Am. Chem. Soc. 2009, 131 (35), 12801–12808.

(19) Pan, J.; Han, J.; Borchers, C. H.; Konermann, L. Characterizing Short-Lived Protein Folding Intermediates by Top-Down Hydrogen Exchange Mass Spectrometry. Anal. Chem. 2010, 82 (20), 8591–8597.

(20) Sterling, H. J.; Williams, E. R. Real-Time Hydrogen/Deuterium Exchange Kinetics via Supercharged Electrospray Ionization Tandem Mass Spectrometry. Anal. Chem. 2010, 82 (21), 9050–9057.

(21) Pan, J.; Han, J.; Borchers, C. H.; Konermann, L. Structure and Dynamics of Small Soluble Aβ(1-40) Oligomers Studied by Top-down Hydrogen Exchange Mass Spectrometry. Biochemistry (Mosc.) 2012, 51 (17), 3694–3703.

(22) Pan, J.; Han, J.; Borchers, C. H.; Konermann, L. Conformer-Specific Hydrogen Exchange Analysis of Aβ(1-42) Oligomers by Top-Down Electron Capture Dissociation Mass Spectrometry. Anal. Chem. 2011, 83 (13), 5386–5393.

(23) Pan, J.; Borchers, C. H. Top-down Mass Spectrometry and Hydrogen/Deuterium Exchange for Comprehensive Structural Characterization of Interferons: Implications for Biosimilars. PROTEOMICS 2014, 14 (10), 1249–1258.

(24) Pan, J.; Zhang, S.; Parker, C. E.; Borchers, C. H. Subzero Temperature Chromatography and Top-down Mass Spectrometry for Protein Higher-Order Structure Characterization: Method Validation and Application to Therapeutic Antibodies. J. Am. Chem. Soc. 2014, 136 (37), 13065–13071.

(25) Going, C. C.; Xia, Z.; Williams, E. R. Real-Time HD Exchange Kinetics of Proteins from Buffered Aqueous Solution with Electrothermal Supercharging and Top-Down Tandem Mass Spectrometry. J. Am. Soc. Mass Spectrom. 2016, 27 (6), 1019–1027.

(26) Li, G.; Reinberg, D. Chromatin Higher-Order Structures and Gene Regulation. Curr. Opin. Genet. Dev. 2011, 21 (2), 175–186.

(27) Woodcock, C. L.; Ghosh, R. P. Chromatin Higher-Order Structure and Dynamics. Cold Spring Harb. Perspect. Biol. 2010, 2 (5), a000596.

(28) Luger, K.; Mäder, A. W.; Richmond, R. K.; Sargent, D. F.; Richmond, T. J. Crystal Structure of the Nucleosome Core Particle at 2.8 Å Resolution. Nature 1997, 389 (6648), 251–260.

(29) Preez, L. L. du; Patterton, H.-G. Secondary Structures of the Core Histone N-Terminal Tails: Their Role in Regulating Chromatin Structure. In Epigenetics: Development and Disease; Kundu, T. K., Ed.; Subcellular Biochemistry; Springer Netherlands, 2013; pp 37–55.

(30) Pepenella, S.; Murphy, K. J.; Hayes, J. J. Intra- and Inter-Nucleosome Interactions of the Core Histone Tail Domains in Higher-Order Chromatin Structure. Chromosoma 2014, 123 (1-2), 3–13.

(31) Black, B. E.; Brock, M. A.; Bédard, S.; Woods, V. L.; Cleveland, D. W. An Epigenetic Mark Generated by the Incorporation of CENP-A into Centromeric Nucleosomes. Proc. Natl. Acad. Sci. 2007, 104 (12), 5008–5013.

(32) Panchenko, T.; Sorensen, T. C.; Woodcock, C. L.; Kan, Z.; Wood, S.; Resch, M. G.; Luger, K.; Englander, S. W.; Hansen, J. C.; Black, B. E. Replacement of Histone H3 with CENP-A Directs Global Nucleosome Array Condensation and Loosening of Nucleosome Superhelical Termini. Proc. Natl. Acad. Sci. 2011, 108 (40), 16588–16593.

(33) Luger, K.; Rechsteiner, T. J.; Richmond, T. J. Preparation of Nucleosome Core Particle from Recombinant Histones. Methods Enzymol. 1999, 304, 3–19.

(34) Sekulic, N.; Bassett, E. A.; Rogers, D. J.; Black, B. E. The Structure of (CENP-A-H4)_2_ Reveals Physical Features That Mark Centromeres. Nature 2010, 467 (7313), 347.

(35) Sekulic, N.; Black, B. E. Preparation of Recombinant Centromeric Nucleosomes and Formation of Complexes with Nonhistone Centromere Proteins. Methods Enzymol. 2016, 573, 67–96.

(36) Kan, Z.-Y.; Mayne, L.; Chetty, P. S.; Englander, S. W. ExMS: Data Analysis for HX-MS Experiments. J. Am. Soc. Mass Spectrom. 2011, 22 (11), 1906–1915.

(37) Lin, S.; Garcia, B. A. Chapter One - Examining Histone Posttranslational Modification Patterns by High-Resolution Mass Spectrometry. In Methods in Enzymology; Carl Wu and C. David Allis, Ed.; Nucleosomes, Histones & Chromatin Part A; Academic Press, 2012; Vol. Volume 512, pp 3–28.

(38) Sidoli, S.; Bhanu, N. V.; Karch, K. R.; Wang, X.; Garcia, B. A. Complete Workflow for Analysis of Histone Post-Translational Modifications Using Bottom-up Mass Spectrometry: From Histone Extraction to Data Analysis. J. Vis. Exp. JoVE 2016, No. 111.

(39) Amon, S.; Trelle, M. B.; Jensen, O. N.; Jørgensen, T. J. D. Spatially Resolved Protein Hydrogen Exchange Measured by Subzero-Cooled Chip-Based Nanoelectrospray Ionization Tandem Mass Spectrometry. Anal. Chem. 2012, 84 (10), 4467–4473.

(40) Wang, L.; Li, D.-Q.; Fu, Y.; Wang, H.-P.; Zhang, J.-F.; Yuan, Z.-F.; Sun, R.-X.; Zeng, R.; He, S.-M.; Gao, W. PFind 2.0: A Software Package for Peptide and Protein Identification via Tandem Mass Spectrometry. Rapid Commun. Mass Spectrom. 2007, 21 (18), 2985–2991.

(41) Perkins, D. N.; Pappin, D. J. C.; Creasy, D. M.; Cottrell, J. S. Probability-Based Protein Identification by Searching Sequence Databases Using Mass Spectrometry Data. ELECTROPHORESIS 1999, 20 (18), 3551–3567.

(42) Black, B. E.; Foltz, D. R.; Chakravarthy, S.; Luger, K.; Woods, V. L.; Cleveland, D. W. Structural Determinants for Generating Centromeric Chromatin. Nature 2004, 430 (6999), 578–582.

(43) Bowman, A.; Ward, R.; El-Mkami, H.; Owen-Hughes, T.; Norman, D. G. Probing the (H3-H4)2 Histone Tetramer Structure Using Pulsed EPR Spectroscopy Combined with Site-Directed Spin Labelling. Nucleic Acids Res. 2010, 38 (2), 695–707.

(44) Spitzer, M.; Wildenhain, J.; Rappsilber, J.; Tyers, M. BoxPlotR: A Web Tool for Generation of Box Plots. Nat. Methods 2014, 11 (2), 121.

(45) Lowary, P. T.; Widom, J. New DNA Sequence Rules for High Affinity Binding to Histone Octamer and Sequence-Directed Nucleosome Positioning. J. Mol. Biol. 1998, 276 (1), 19–42.

(46) Zheng, C.; Lu, X.; Hansen, J. C.; Hayes, J. J. Salt-Dependent Intra- and Internucleosomal Interactions of the H3 Tail Domain in a Model Oligonucleosomal Array. J. Biol. Chem. 2005, 280 (39), 33552–33557.

(47) Ebralidse, K. K.; Grachev, S. A.; Mirzabekov, A. D. A Highly Basic Histone H4 Domain Bound to the Sharply Bent Region of Nucleosomal DNA. Nature 1988, 331 (6154), 365.

(48) Kalashnikova, A. A.; Porter-Goff, M. E.; Muthurajan, U. M.; Luger, K.; Hansen, J. C. The Role of the Nucleosome Acidic Patch in Modulating Higher Order Chromatin Structure. J. R. Soc. Interface 2013, 10 (82), 20121022.

(49) Arya, G.; Schlick, T. Role of Histone Tails in Chromatin Folding Revealed by a Mesoscopic Oligonucleosome Model. Proc. Natl. Acad. Sci. U. S. A. 2006, 103 (44), 16236–16241.

(50) Kan, P.-Y.; Caterino, T. L.; Hayes, J. J. The H4 Tail Domain Participates in Intra- and Internucleosome Interactions with Protein and DNA during Folding and Oligomerization of Nucleosome Arrays. Mol. Cell. Biol. 2009, 29 (2), 538–546.

(51) D’Arcy, S.; Martin, K. W.; Panchenko, T.; Chen, X.; Bergeron, S.; Stargell, L. A.; Black, B. E.; Luger, K. Chaperone Nap1 Shields Histone Surfaces Used in a Nucleosome and Can Put H2A-H2B in an Unconventional Tetrameric Form. Mol. Cell 2013, 51 (5), 662–677.

(52) Lee, K.-M.; Hayes, J. J. The N-Terminal Tail of Histone H2A Binds to Two Distinct Sites within the Nucleosome Core. Proc. Natl. Acad. Sci. 1997, 94 (17), 8959–8964.

